# Flower-like patterns in multi-species biofilms

**DOI:** 10.1101/550996

**Authors:** Liyang Xiong, Yuansheng Cao, Robert Cooper, Wouter-Jan Rappel, Jeff Hasty, Lev Tsimring

## Abstract

Diverse interactions among species within bacterial biofilms often lead to intricate spatiotemporal dynamics. The spatial structure of biofilms can determine growth and survival of different species, but the mechanisms driving formation of this structure are not fully understood. Here, we describe the emergence of complex structures in a biofilm grown from mixtures of motile and non-motile bacterial species on a soft agar surface. Time-lapse imaging shows that non-motile bacteria “hitchhike” on the motile bacteria as the latter migrate outward. The non-motile bacteria accumulate at the boundary of the colony and trigger an instability that leaves behind striking flower-like patterns. The mechanism of the front instability governing this pattern formation is elucidated by a mathematical model for the frictional motion of the colony interface, with friction depending on the local concentration of the nonmotile species along the interface. A more elaborate two-dimensional phase-field model that explicitly accounts for the interplay between growth, mechanical stress from the motile species, and friction provided by the non-motile species, fully reproduces the observed flower-like patterns.

## Introduction

Microbial communities inhabit every ecosystem on Earth, from soil to hydrothermal vents to plants to the human gut [1–3]. They often form dense biofilms, whose structures are shaped by biological, chemical, and physical factors [4–6]. In the wild, most biofilms are comprised of multiple bacterial strains. They feature a diverse repertoire of social interactions, including cooperation [7, 8], competition [9], and predation [10]. Bacteria often signal, sense, and respond to each other through secondary metabolites [11] or antibiotic compounds [12], and co-cultures can even exhibit different motility from either species on its own [13]. These interactions may lead to the emergence of complex spatial structures, which can have a profound effect on bacteria survival and function, and promote biodiversity by optimizing the division of labor within the biofilm [14]. Spatial structure can also enhance horizontal gene transfer among different species [15].

In addition to biochemical interactions, mechanical forces also play an important role in shaping the bacterial community structures. In dense colonies, bacteria push against each other due to growth and motility. Bacteria can exploit these mechanical interactions to adapt to the environment. For example, mechanical stresses generate buckling in *Bacillus subtilis* biofilms to optimize nutrient transport and consumption [16–18]. Although the role of mechanical interactions in single-species colonies has been studied previously [19–22], dynamics of multi-species communities driven by mechanical forces have received much less attention. Since bacterial strains can have significant differences in their growth and motility characteristics, one can expect the mechanical stress distribution to be highly heterogeneous, which in turn can result in a complex spatial structure of the colony.

Here we study the interactions between bacterial species with significantly different biological and physical properties. We choose *Acinetobacter baylyi*, a gram-negative bacterium that easily moves on soft surfaces using twitching motility [23–25], and an *Escherichia coli* strain that is almost non-motile on soft agar. Meanwhile, wild-type *A. baylyi* possesses a Type VI Secretion System (T6SS) that enables them to kill other bacteria (including *E. coli*) on direct contact [15, 26]. We found that when these two strains are mixed together and inoculated on an agar surface, growing biofilms develop intricate flower-like structures that are absent when either species is grown by itself.

To shed light on the mechanism behind this intricate pattern formation, we tested whether biological cell-cell communication or mechanical interaction between strains with different motilities played the key role. Experiments with *A. baylyi* mutants lacking T6SS showed that the pattern formation did not rely on this mechanism. On the other hand, genetically impairing *A. baylyi* motility eliminated the patterns entirely. We also showed that agar concentration that affects cell motility, also played a key role in pattern formation. These findings suggested that the mechanical interactions between species were indeed primarily responsible for the pattern formation.

We then performed theoretical and computational analysis of two models: a geometrical model of the colony boundary motion and a 2D phase-field model of the entire colony to describe the mechanical interaction between two species. Our results show that cell motility differences are sufficient to explain the emerging patterns. Since the structures require only differential motility, this mechanism of pattern formation may be broadly generalizable to other mixed-species biofilms.

## Results

### Flower-like patterns in mixtures of *A. baylyi* and *E. coli* on nutrient-rich soft agar

We inoculated a mixture of *E. coli* and *A. baylyi* cells with an initial density ratio of 10:1 at the center of a Petri dish filled with soft LB agar (0.5% agar). To distinguish the two strains, we labeled *E. coli* with constitutively expressed mTFP. After growing at 37°C for 3 days, this colony developed an intricate flower-like pattern (Fig. 1A). To see how such patterns form, we tracked the colony growth with time-lapse imaging (Fig. 1B, Movie S1). Up to 8 hours after inoculation, the expanding colony remained nearly uniform and circular. Then the colony front began visibly undulate. As the colony expanded further, the undulations grew and formed cusps that in turn would leave behind tracks (or “branches”). These branches then merged, following the movement of cusps along the interface as the colony continued to expand. The branches were visible even in bright-field imaging, but they were also bright in the teal fluorescence channel, indicating that branches predominantly consisted of *E. coli* cells (Fig. S1).

**Figure 1:**
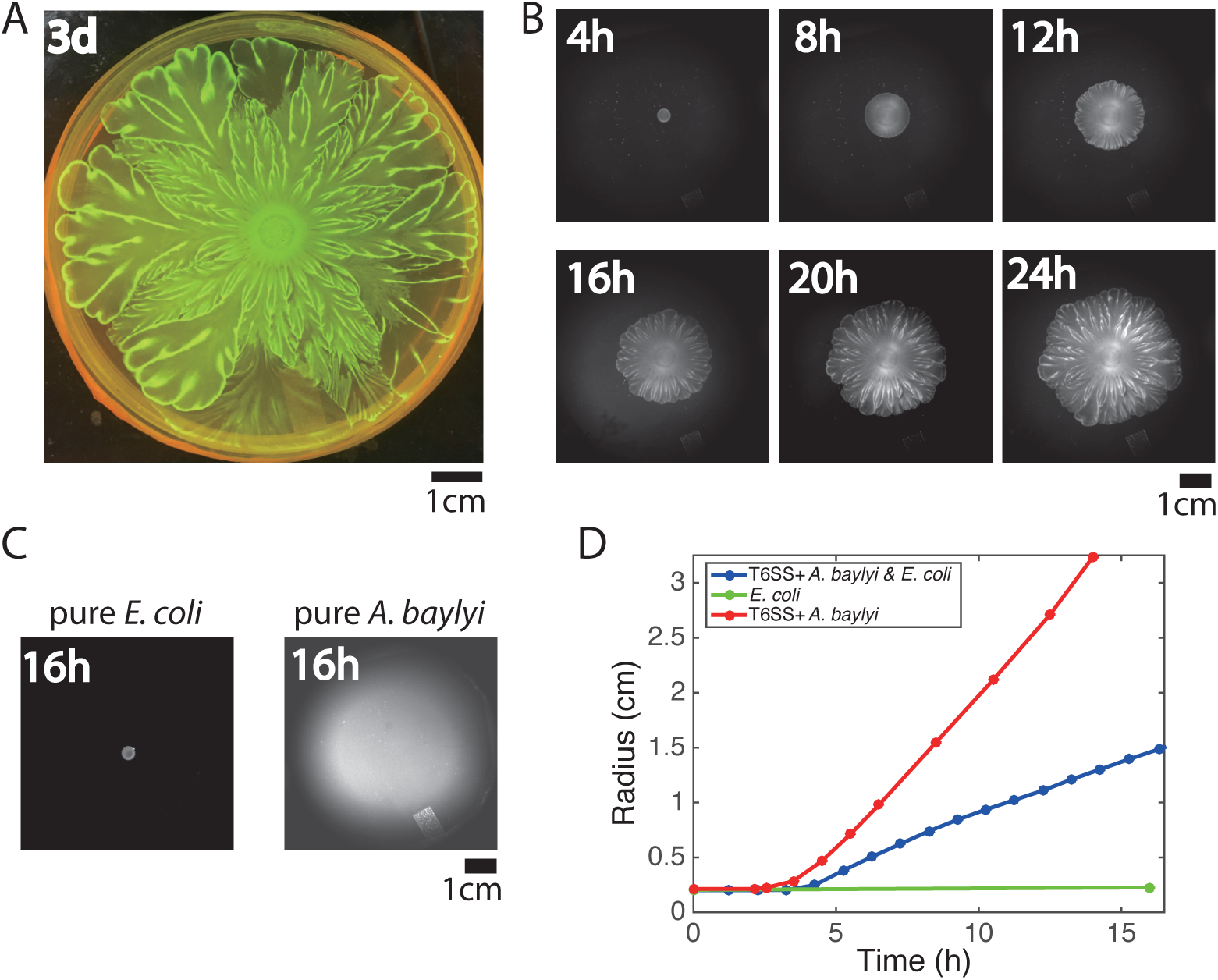
Flower-like patterns in mixtures of *E. coli* and *A. baylyi*. (A) The pattern after 3 days of growth on a 0.5% LB agar surface. (B) Time-lapse bright-field images of the developing pattern. (C) Pure *E. coli* and pure *A. baylyi* colonies show no patterns. (D) Radius of the colony vs time for pure *E. coli* (green), pure *A. baylyi* (red), and the mixture of *E. coli* and *A. baylyi* (blue).

To test whether these flower-like patterns originate from interactions between the two species, we grew each species separately on the same 0.5% LB agar surface. The *E. coli* motility on agar is small, and the colony size remained relatively unchanged after 16 hours of growth (Fig. 1C, left). After the same time, a colony of highly motile *A. baylyi* reached the edge of the plate (Fig. 1C, right). In both cases, no patterns emerged, which proves that the flower-like pattern formation is a result of inter-species interaction. We measured the sizes of mixed, pure *E. coli* and pure *A. baylyi* colonies at different times after inoculation (Fig. 1D). The expansion speed of mixed colonies falls between those of pure *A. baylyi* and pure *E. coli* colonies, and the speed did not change much once the colonies began expanding.

### *E. coli* destabilize colony front by hindering *A. baylyi* expansion

To observe the pattern formation at higher resolution, we modified the experimental setup to fit under a fluorescence microscope (see Methods). After 24 hours of growth, a droplet of 1:1 mixture of *E. coli* (expressing mTFP) and *A. baylyi* (expressing mCherry) grew into a clearly-visible flower-like pattern (Fig. 2A). By zooming in on the front of the expanding colony, we were able to track the formation and merging of branches that give rise to the flower-like structure of the patterns (Fig. 2B, Movie S2). While *A. baylyi* managed to kill most of *E. coli* via T6SS within the inoculum, a significant number of *E. coli* managed to survive at the periphery where they were not in direct contact with *A. baylyi*. *E. coli* also has a higher growth rate (1.53 ± 0.11 h^−1^, *n* = 3) than *A. baylyi* (1.13 ± 0.03 h^−1^, *n* = 3), so by the time the colony began to expand, *E. coli* cells had already grown near the colony boundary which resulted in a band of *E. coli* around the expanding colony of mostly *A. baylyi* (Fig. 2B, 11h).

**Figure 2:**
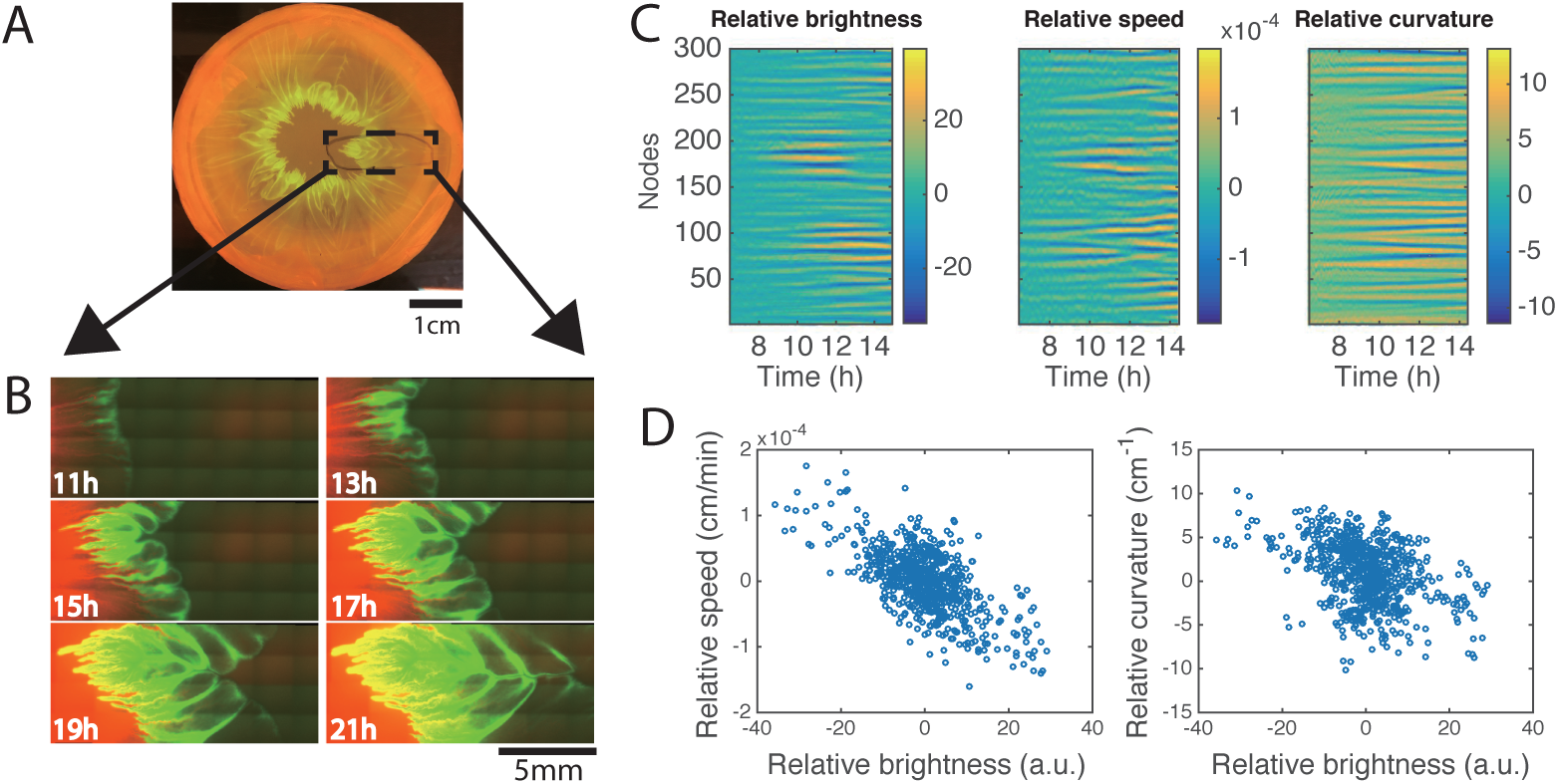
Development of branches in a growing pattern. (A) The whole colony in a Petri dish after one day. (B) Time-lapse microscopic images of the front propagation leading to branch formation and merging. (C) Kymographs of relative brightness, speed and curvature along the colony boundary. (D) Scatter plots for relative brightness and speed (left) and relative brightness and curvature (right). Each circle corresponds to one virtual tracking node at one time point.

As the colony kept expanding, in regions with more *E. coli* cells near the front, the expansion was slower, so the interface began to curve inward (Fig. 2B, 13h). As the undulations grew bigger, the *E. coli* in the regions lagging behind became more concentrated, thus slowing down the local front advance even more. Eventually, the front folded onto itself near these stagnant regions and formed narrow “branches” that continued to grow outward with the expanding colony front (Fig. 2B, 15h, 17h). Later, the front with the branches folded again, and the previous branches merged inside the new fold (Fig. 2B, 19h, 21h). Since *E. coli* continued to grow at the expanding colony front, new undulations and branches constantly appeared, and eventually a macroscopic, flower-like pattern of growing and converging branches formed.

To quantify the effect of local *E. coli* concentration on the biofilm expansion, we analyzed the time-lapse images in Fig. 1B (see Methods). We adapted a boundary tracking program for eukaryotic cells [27] to track the boundary of the bacterial colony. The colony boundary was parameterized by 300 virtual “nodes” connected by springs [28]. For each node, we measured local brightness (a proxy for *E. coli* concentration), speed and curvature. To offset the non-uniformity of the illumination and the overall change in speed and curvature for a growing colony, we detrended the data. The kymographs of these quantities for each node are shown in Fig. 2C. Then we compute cross-correlations between these quantities within the time window when the pattern began to form (about 9.5 - 11.5 hours after inoculation). As shown in Fig. 2D (left), the brightness and extension speed show an anti-correlation (Pearson coefficient *ρ* = −0.66). This result confirms that higher *E. coli* density slows down the front propagation. Variations in the front speed lead to variations of the local curvature, and the scatter plot between brightness and curvature indeed shows significant anti-correlation (Fig. 2D right, Pearson coefficient *ρ* = −0.41).

### Robustness of flower-like patterns to perturbations

First, we explored the effect of the initial *A. baylyi*:*E. coli* (A:E) density ratio on the resulting pattern. We varied the ratio of *A. baylyi* to *E. coli* in the inoculum while maintaining the same total density of bacteria. We found that when the starting ratios are low (A:E= 1:100 and 1:10), flower-like patterns emerged, while at high ratios (10:1 and 100:1) the *E. coli* were completely eliminated and no patterns formed (Fig. 3A). At the intermediate ratio 1:1, *A. baylyi* dominated significantly at the center of the colony by killing *E. coli*, but the flower-like structure still developed at the colony periphery.

**Figure 3:**
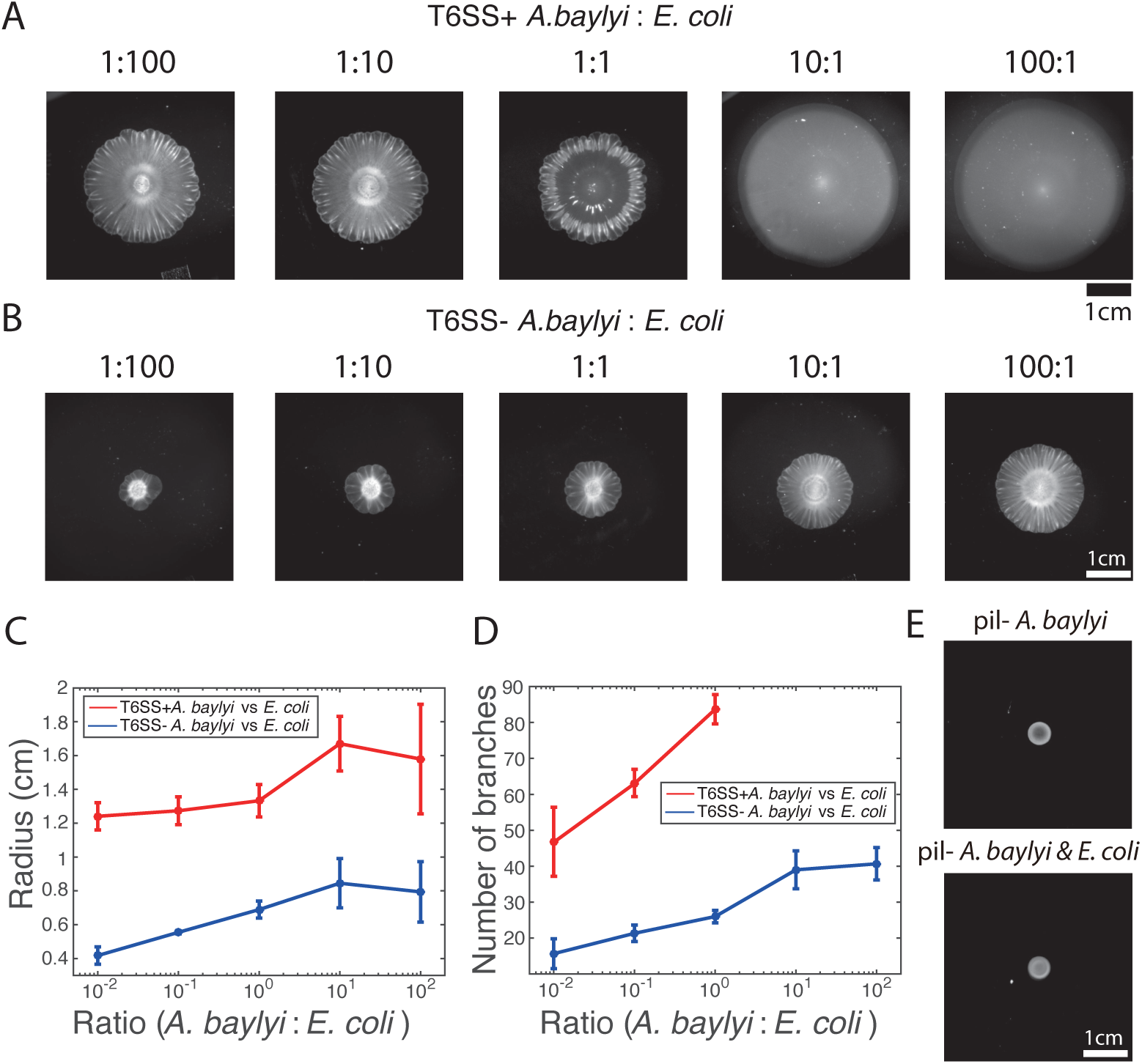
Pattern formation requires *A. baylyi* motility, but not killing. (A,B) Bright-field snapshots of colonies of T6SS^+^ (A) and T6SS^−^ (B) *A. baylyi* with *E. coli* 16 hours after inoculations at different initial density ratios. (C) The average colony radius vs density ratios. (D) Number of branches at the onset of front instability vs density ratios. (E) Colonies of pure pil^−^ T6SS^+^ *A. baylyi* and the mixture of pil^−^ T6SS^+^ *A. baylyi* and *E. coli* 16 hours after inoculation.

Second, we wondered whether T6SS-dependent killing played a role in the formation of these patterns when *E. coli* were not completely eliminated. We tested this by knocking out T6SS in *A. baylyi* (see Methods for details). The growth rate of T6SS^−^ *A. baylyi* (1.09 ± 0.01 h^−1^, *n* = 3) was not significantly different from the wild type while their motility was slightly lower as determined by colony expansion rate. Still, it was still much higher than *E. coli* (Fig. S2). We inoculated mixtures of T6SS^−^ *A. baylyi* and *E. coli* with different initial ratios on 0.75% LB agar, and observed that the colony still formed an outer ring of *E. coli* (Fig. S3) and subsequently developed front instability, branches of *E. coli*, and a flower-like pattern in all cases (Fig. 3B). The only qualitative difference between the T6SS^−^ and T6SS^+^ cases was that in the nonkilling case *E. coli* remained at a high concentration within the area of the initial inoculum. We measured the average radius of the colonies with different initial density ratios (Fig. 3C). In the case of mixture of T6SS^−^ *A. baylyi* and *E. coli*, the more *E. coli* in the inoculum, the slower the colony expanded, which is consistent with our hypothesis that *E. coli* hinders the overall colony expansion. However, the trend is not as significant for T6SS^+^ case because, as we reason, T6SS^+^*A. baylyi* kill most *E. coli* at the early stage, which increases and stabilizes the effective A:E ratio. We also counted the number of branches as they first emerged, and found more branches in colonies seeded with less *E. coli* (Fig. 3D). In general, the overall structure of the patterns remained unchanged in the mixture of T6SS^−^ *A. baylyi* and *E. coli*. Thus, we concluded that the T6SS did not play a major role in the formation of flower-like patterns.

Third, the fact that two-species colonies expanded much more quickly than pure *E. coli* colonies strongly suggested that the high motility of *A. baylyi* is primarily responsible for the colony expansion. To test this hypothesis, we knocked out the *pilTU* locus of T6SS^+^ *A. baylyi*, which is required for the pilus-based twitching motility of *A. baylyi* [25, 29]. As expected, colonies of *pilTU*^−^ *A. baylyi* cells did not expand significantly (Fig. 3E, top) and did not form branching patterns when mixed with *E. coli* cells (Fig. 3E, bottom). This demonstrates that the high *A. baylyi* motility plays a crucial role in the flower-like pattern formation.

### Pattern-forming instability originates at the colony interface

Experiments showed that the formation of flower-like patterns appears to be preceded and caused by growing undulations of the colony front, where *E. coli* cells concentrate and locally slow expansion. To mechanistically understand how a ring of low-motility bacteria surrounding an expanding core of highly-motile bacteria can create such patterns, we turned to mathematical modeling. We adapted a one-dimensional “geometrical” model of front dynamics [30,31] that casts the motion of the interface in natural, reference-frame independent variables of curvature *κ* and metric *g* as a function of its arclength *s* and time *t*:

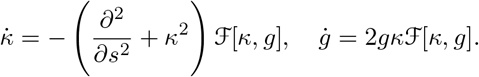

In the overdamped limit, the velocity functional 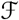 is proportional to the balance of a constant outward pressure force *F*_0_ due to *A. baylyi* motility, surface tension *F*_*s*_ proportional to the interface curvature, and the resistance (friction) force *F*_*r*_that is assumed to be proportional to the local concentration of *E. coli* on the interface *c*(*s, t*) (Fig. 4A). For simplicity we ignore *E. coli* growth and leakage from the boundary in the interior and assume that the local concentration of *E. coli* is only changed by stretching or contraction of the interface, therefore *c* is taken to be inversely proportional to the metric *g*. A straightforward linear stability analysis demonstrates that the interface is indeed unstable to a broad spectrum of initial perturbations (for more details see Supporting Information).

**Figure 4:**
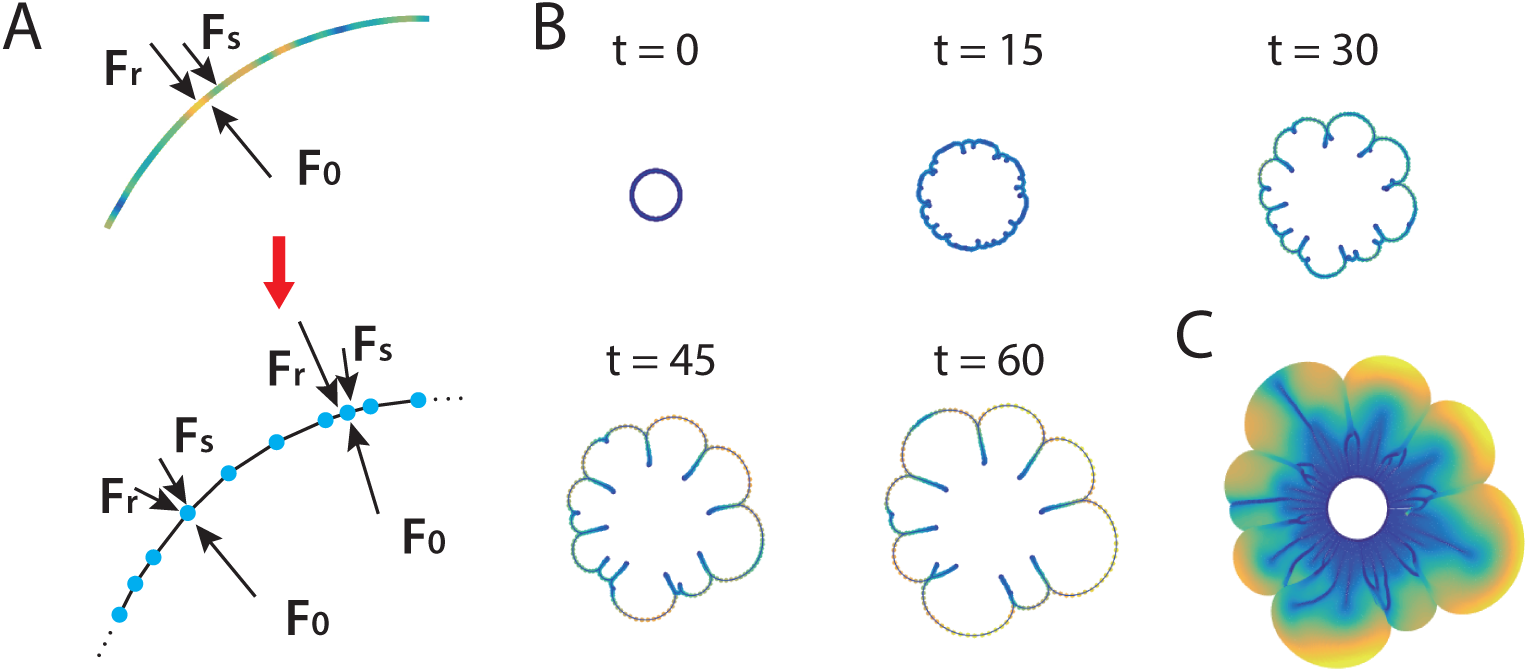
Discrete interface model. (A) Sketches of the continuum and discrete interface models. (B) Snapshots of the interface in discrete interface model for a sample simulation with parameters listed in Supporting Information. The colors of the nodes correspond to the distance between node and its neighbors. (C) “Fossil record” of *E. coli* densitiy on the moving interface.

To simulate the interface dynamics beyond the linear regime, we also constructed a discrete model of the continuous interface by replacing it with a closed chain of nodes connected by straight links (Fig. 4A bottom). Each node carries a fixed amount of *E. coli*, so the local density of nodes per unit length of the interface corresponds to the local density of *E. coli*. Nodes are driven by a constant outwards expansion force *F*_0_, surface tension, and a friction force that is proportional to the window-weighted average density of nodes per unit length. Additionally, we introduced short-range repulsive forces among nodes and between nodes and links, to prevent self-crossing of the interface. Detailed description of this model is also given in Supporting Information.

As an initial condition, we assumed that the chain forms a circle with nodes slightly perturbed from equidistant positions. Figure 4B shows time-lapse snapshots of the interface in a sample simulation (also see Movie S3). Fig. 4C shows the aggregate image of the interface during the colony expansion, with the color of a point corresponding to density of nodes when the interface passed through that point (also see Movie S4). Assuming that a fixed fraction of *E. coli* is left behind the interface, this interface “fossil record” should roughly correspond to the density of *E. coli* inside the colony. At the beginning, the interface remains nearly circular, but initial perturbations quickly grow as the colony expands, producing large front undulations. Regions with lower node density expand more quickly because they experience less friction, and this expansion stretches the chain and further reduces the node density per unit length, creating a positive feedback loop. Concave regions, on the contrary, accumulate nodes and thus move outward more slowly. Eventually, cusps are formed in these lagging regions that have very high node density and therefore move very slowly, if at all. The regions on both sides of the cusp continue to expand toward each other and eventually “collide”. After collision they form “double-layers” that remain nearly static and only increase in length as the overall interface expands further. Thus, “branches” with high concentration of *E. coli* form. As the front continues to expand, the interface already containing branches continues to undulate and form new cusps. This causes the earlier branches to merge, similar to what we observed in experiments (Fig. 2B). These simulation results suggest that indeed branch formation and merging can be explained by mechanics of a resistive ring surrounding a colony, which is stretched by the colony expansion. However, since this model neglects *E. coli* growth, the average density of nodes per unit length gradually decays, and eventually, the front instability ceases, in divergence with experimental results. To account for cell growth as well as for the diffusive leakage of *E. coli* from the interface into the bulk of the expanding colony, we developed a more elaborate 2D model of the growing multi-species biofilm.

### Phase-field model of flower-like pattern formation

Our two-dimensional, multi-component model is conceptually similar to the phase-field models used for description of eukaryotic cell motility and migration [32–34] (Fig. 5A). It is based on three PDEs for the densities of *A. baylyi ρ*_*A*_, *E. coli ρ*_*E*_, and the phase field *ϕ* that changes continuously from 1 inside the colony to 0 outside (see Supporting Information for detailed formulation of the model). The latter is introduced to avoid computational difficulties of dealing with the sharp colony interface. All three components are diffused and advected by the velocity field that is generated by a combination of stress due to cell growth and motility, viscosity, and bottom friction that is proportional to the local *E. coli* density.

**Figure 5:**
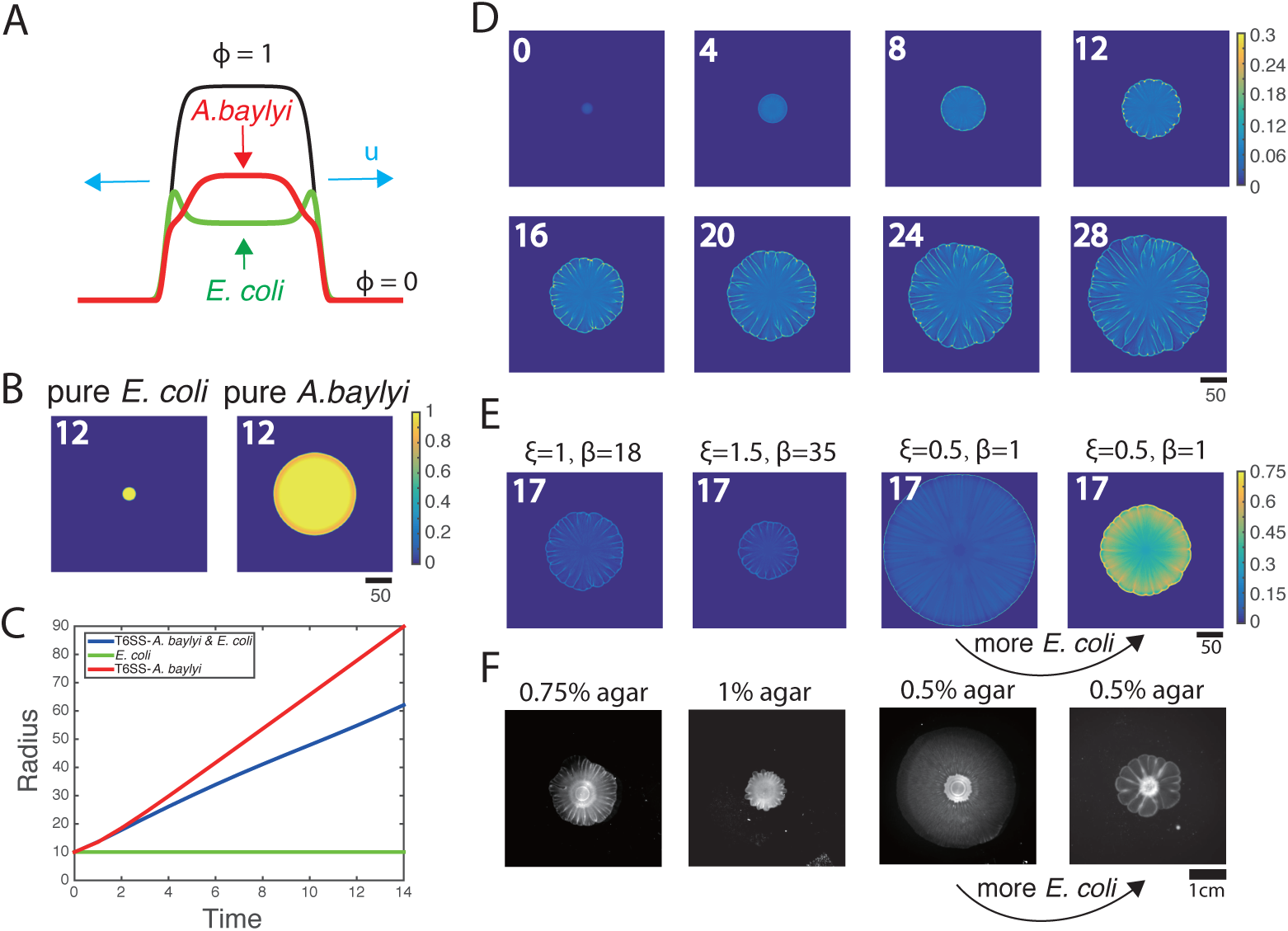
Phase-field model simulations of two-species colony growth. (A) Illustration of the model. (B) Snapshots of the colonies of pure *E. coli* and pure *A. baylyi* at *t* = 12. A colony of *E. coli* expanded only slightly, while a pure colony of *A. baylyi* expanded quickly, but remained circular. (C) Colony radius vs time for the mixed and single-species colonies. (D) Several snapshots of *E. coli* density during the growth of a mixed colony in simulations. (E) Colony snapshots at time *t* = 17 in simulations using different friction parameters. For larger friction, the colony grew slower, but still featured flow-like patterns. For smaller friction, the colony expanded more quickly, but patterns eventually disappeared. However, increasing the initial concentration of *E. coli* at low friction coefficients restored patterning. (F) Experimental snapshots with different agar concentrations 16 hours after inoculation: similar phenomenology observed.

When we initialized the model with small circular domains of either pure *E. coli* or *A. baylyi*, the colony boundaries remained circular, and no patterns emerged (Fig. 5B). Consistent with the experiments, the *E. coli* colony only slightly expanded, while the *A. baylyi* colony expanded rapidly (Fig. 5C). When we initialized the model with a mixture of *A. baylyi* and *E. coli*, the colony grew at an intermediate speed (Fig. 5C), as in the experiments (Fig. 1D). The mixed colony simulations also exhibited front instability leading to formation of branches of *E. coli* (Fig. 5D, the snapshots of *A. baylyi* are shown in Fig. S4, also see Movie S5). As the colony grew, the branches merged and expanded, and a flower-like pattern developed.

Agar concentration is known to have a strong effect on the motility of bacteria [23] and their adhesion to the agar surface [35], so we reasoned that in our phase-field model changing agar concentration should be reflected in changing friction parameters. The frictional force in our model consists of two contributions: a small basal friction (characterized by parameter *ξ*) and stronger contribution proportional to the local *E. coli* concentration with coefficient *β*. Thus, to mimic different agar concentrations, we varied both *ξ* and *β*. The leftmost panel in Fig. 5E shows the colony snapshots at *t* = 17 for the same parameter values as the time-lapse sequence in Fig. 5D. The next panel corresponds to larger *ξ* and *β* (presumably, higher agar concentration), where as expected, the colony expanded slower. The third panel shows the snapshot for smaller *ξ* and *β* (lower agar concentration), in which case the colony expands fast, but no patterns emerge. However, for the same low *ξ* and *β*, when we started a simulation from 10x higher *E. coli* density, the friction provided by *E. coli* increased, and patterning re-emerged (Fig. 5E, fourth panel).

These numerical predictions were fully validated by experiments in which we varied the agar concentration and the initial density ratio of *E. coli* and T6SS^−^ *A. baylyi*. The leftmost panel in Fig. 5F shows the snapshot of the colony started from 1:1 mixture after 16 hours of growth on 0.75% agar surface. When we increased the agar concentration to 1% (Fig. 5F, second panel), the colony expanded slower but the flower-like pattern emerged. Conversely, for low agar concentration (0.5%), colony grew fast but patterns were completely eliminated (Fig. 5F, third panel). However, for the same 0.5% agar concentration but A:E=1:100 initial density ratio, the flower-like pattern formation was rescued (Fig. 5F, fourth panel).

## Discussion

Motility plays a key role in the spread of dense bacterial colonies. In this paper, we studied the structure of growing biofilms comprised of two bacterial species, *E. coli* and *A. baylyi*, with very different motilities. Not only did the highly-motile species (*A. baylyi*) accelerate the spread of the slow species (*E. coli*), but the structure of the expanding colony quickly became highly heterogeneous and eventually produced very intricate, flower-like patterns.

Pattern formation in growing colonies of single bacterial species has been studied extensively [36–39], and branching patterns were often found in these experiments. The emergence of these patterns is usually driven by nutrient limitation and ensuing chemotaxis, with agar concentration also having a strong effect on their morphology. For example, colonies expand homogeneously on soft agar rich with nutrients, but under nutrient limitation and in semi-solid agar, complex patterns emerge [37,39]. In our system, however, we used rich LB media, and single-species colonies in the same conditions did not produce patterns. This suggested that the mechanism of pattern formation here was different. We also found no significant differences in pattern formation with T6SS^+^ and T6SS^−^ strains of *A. baylyi*. In fact, we did not observe noticeable killing of *E. coli* by T6SS^+^ *A. baylyi* after a short initial period (Movie S2). We believe that an extracellular matrix may have played a role here, as recent studies showed that it protected bacteria from T6SS attacks from other species [40, 41]. Overall, our experiments and modeling provided strong evidence in favor of the mechanical nature of the pattern-forming instability, arising from the interplay between outward pressure generated by the growth and high motility of *A. baylyi*, and the friction provided by sessile *E. coli* that adhere to the agar surface.

Ecologically, one of the primary challenges for any species is to maximize its geographic dispersal. Motility enables bacteria to escape from local stresses, move to locations with more nutrients, or invade host tissue [23]. However, motility, especially on hard surfaces, requires additional gene expression which could be a metabolic burden [21]. So some bacteria take advantage of other species with larger motility to colonize new niches. For example, by hitchhiking on zooplankton, water-borne bacteria can reach places that are otherwise inaccessible for them due to density gradients [42]. Non-motile Staphylococcal species hitchhike on swimming bacteria such as *Pseudomonas aeruginosa* [43]. In other studies, motile swarming *Paenibacillus vortex* was shown to transport non-motile *Xanthomonas perforans* [44] or *E. coli* [45] on agar surfaces. In our system, *A. baylyi* cells move by twitching instead of swarming, and our results suggest that slow-moving bacteria might take advantage of fast-moving twitching species by hitchhiking, or “surfing” along the expanding boundary, and thus spread farther. This can be seen clearly from the experiment in which *E. coli* and *A. baylyi* were inoculated separately at a small distance on agar surface (Movie S6). *A. baylyi* colony expanded and pushed *E. coli* to places where *E. coli* alone could not reach. Although *E. coli* and *A. baylyi* may not necessarily find themselves in the same ecological niche, bacteria with different motilities are ubiquitous in the environment [23]. Therefore, the mechanisms of codependent motility and pattern formation described here are likely to be broadly applicable in natural habitats or even have implications in the transmission of pathogenic microbes. For example, *Acinetobacter baumannii*, an increasing threat in hospitals due to mutil-drug resistance [46], is closely related to *A. baylyi* [15, 47] and also has twitching motility [48, 49], and thus other microbes which coexist with *A. baumannii* might also take advantage of its motility to spread.

## Materials and Methods

### Strains

We used *E. coli* MG1655 and *A. baylyi* ADP1 (ATCC #33305). The *E. coli* strain carried a plasmid that constitutively expressed mTFP and a kanamycin resistance gene. *A. baylyi* had a kanamycin resistance gene and the mCherry gene integrated in the genome. We also constructed a T6SS^−^ *A. baylyi* (Δhcp) mutant by first fusing the tetracycline resistance marker from pTKS/CS to approximately 400 bp homology arms amplified from either side of hcp (ACIAD2689) in the *A. baylyi* genome, and mixing the donor oligo with naturally competent *A. baylyi*. The *pilTU*^−^ strain was constructed similarly to delete the genes ACIAD0911-0912.

### Culture conditions and image capturing

*E. coli* and *A. baylyi* cells were taken from −0°C glycerol stocks, inoculated in LB with appropriate antibiotics (kanamycin for *E. coli* and T6SS^+^ *A. baylyi*, tetracycline for T6SS^−^ *A. baylyi*) and grown at 37°C separately. When their OD600 reached about 0.3, both *E. coli* and *A. baylyi* were concentrated to OD=1, still separately. They were then mixed at specified volume ratios, and 3 *μL* was inoculated on the surface of 10 mL LB agar in the center of an 8.5 cm Petri dish. The plate was incubated at 37°C. The images were taken using a custom “milliscope” fluorescence imaging device unless indicated otherwise.

When the colony development was to be observed under a microscope, a 5.5 cm Petri dish was used with 15 mL 1% base agar (without LB) and top 10 mL LB agar (1% agar). After the cell culture was inoculated and dried, it was put on the stage of an inverted, epifluorescence microscope (Nikon TI2). The magnification was 4X. Fluorescent images were acquired using a 4X objective and a Photometrics CoolSnap cooled CCD camera in a 37°C chamber. The microscope and accessories were controlled using the Nikon Elements software.

The bacteria growth rates were measured in a Tecan plate reader.

### Colony tracking

We adapted the method and the MATLAB^™^ code from [27] to track the colony boundary. The bright-field images were first segmented to identify the colony using an active contour method [50]. The segmentation result is illustrated in Movie S7. Then the colony boundary pixels were interpolated by a closed cubic spline and the boundary was parameterized by 300 virtual nodes, which were evolved in time as a coupled spring system (Fig. S5) [28]. For each node, three quantities were measured: brightness, extension speed and curvature. Brightness at each node was defined as the median of the neighboring pixels assigned to each node (see [27]). Extension speed was computed by the displacement of a node from frame *t* to frame *t* + 10. Curvature was calculated by taking derivatives of the spline contour. Then the time series of these quantities were detrended and smoothed using FFT. An example of these quantities for all nodes at a particular time point is shown in Fig. S6. In Fig. 2D, we sampled 7 time points with 20 min interval from 9.5 h to 11.5 h and for each time point we plotted 100 nodes.

### Mathematical models

Detailed description of the two models is given in Supporting Information.

## Acknowledgments

We thank Megan Dueck for building the custom “milliscope” used in our study and Kit Pogliano lab for providing the original *E. coli* strain. This work was supported by the National Institutes of Health (grant R01-GM069811), the National Science Foundation (grant PHY-1707637), San Diego Center for Systems Biology (NIH grant P50-GM085764) and the DOD Office of Naval Research (grant N00014-16-1-2093).

## Author Contributions

L.X. and L.T. designed research; L.X., R.C. did experiments and analyzed data; L.X., Y.C. and L.T. did modeling; L.X, Y.C., R.C., W.R., J.H. and L.T. analyzed results and wrote the paper.

## Author Declaration

The authors declare no conflict of interest.

