## Supplementary Text for "Flower-like patterns in multi-species biofilms"

### Supporting Information
### Flower-like patterns in multi-species biofilms

Liyang Xiong[a,b], Yuansheng Cao[a], Robert Cooper[b], Wouter-Jan Rappel[a], Jeff Hasty[b,c,d,e], and Lev Tsimring[b,c*]

[a]Department of Physics, University of California, San Diego, La Jolla, CA 92093-0374
[b]BioCircuits Institute, University of California, San Diego, La Jolla, CA 92093-0328
[c]The San Diego Center for Systems Biology, La Jolla, CA 92093-0688
[d]Molecular Biology Section, Division of Biological Sciences, University of California, San Diego, La Jolla, CA 92093
[e]Department of Bioengineering, University of California, San Diego, La Jolla, CA 92093

#### Interface model

##### Continuous interface dynamics

To describe the motion of the interface separating the growing bacterial colony from the environment, we can use the framework originally proposed by Brower et al. [1, 2] for solidification patterns. The 1D interface (a closed line) at time $t$ is specified by the position vector $\mathbf{x}(t, \sigma)$ where $0 < \sigma < 1$ is the variable parametrizing the interface such that $\mathbf{x}(t, 0) = \mathbf{x}(t, 1)$. Using the "orthogonal gauge" assumption that the velocity $d\mathbf{x}/dt$ is orthogonal to the tangent vector $\boldsymbol{\tau} = \partial\mathbf{x}/\partial\sigma$, the equation of motion for the interface can be written in the general form

$$\frac{d\mathbf{x}}{dt} = \hat{\mathbf{n}}\mathcal{F}(\mathbf{x}, \partial\mathbf{x}/\partial\sigma, ...) \tag{S1}$$

where $\hat{\mathbf{n}}$ is the unit normal to the interface at $\mathbf{x}$ (perpendicular to $\boldsymbol{\tau}$) and $\mathcal{F}$ is the velocity functional that generally may depend on the overall interface position and other parameters. As Brower et al. [2] demonstrated, this equation can be transformed to the reference-frame independent *local* equations of motion for the local curvature $\kappa$ and the curve metric $g = \boldsymbol{\tau} \cdot \boldsymbol{\tau}$ as a function of arclength $s$ and time $t$:

$$\dot{\kappa} = -\left(\frac{\partial^2}{\partial s^2} + \kappa^2\right)\mathcal{F} \tag{S2}$$

$$\dot{g} = 2g\kappa\mathcal{F} \tag{S3}$$

Here the arclength is given by

$$s = \int_0^\sigma \sqrt{g(\sigma')}d\sigma' \tag{S4}$$

and the curvature is defined by

$$\kappa = -\hat{\mathbf{n}} \cdot \frac{\partial^2\mathbf{x}}{\partial s^2} \tag{S5}$$

Now we need to specify the velocity functional $\mathcal{F}$ for our system in which a growing colony is surrounded by the thin band of highly frictional *E. coli* that hinders the colony expansion. Thus, we assume that $\mathcal{F}$ depends only on the local curvature $\kappa$ and the local concentration of *E. coli* on the interface, $c$. This assumption will be violated if/when the interface will develop large folds and will attempt to "collide" with each other, then non-local terms in $\mathcal{F}$ become essential. We confine our continuous description here to sufficiently early times before this non-local interaction occurs. Under the additional simplifying assumption that the total amount of *E. coli* on the interface is conserved and neglecting their diffusion along the interface, the local concentration of *E. coli* will be inversely proportional to the metric $g$, $c = c_0/g$. In reality, of course, *E. coli* also grows and is left behind in the bulk of the colony, but we assume that in a quasi-stationary regime these two processes approximately balance each other. Thus, the closed-form model for the interface expansion has the following form

$$\dot{\kappa} = -\left(\frac{\partial^2}{\partial s^2} + \kappa^2\right)\mathcal{F}(\kappa, c_0/g) \tag{S6}$$

$$\dot{g} = 2g\kappa\mathcal{F}(\kappa, c_0/g) \tag{S7}$$

For specificity, we assume that $\mathcal{F}$ has the following simple form:

$$\mathcal{F} = F_0 - \gamma\kappa - \alpha c \tag{S8}$$

where $F_0$ is the maximal expansion velocity and $c$ is the concentration of *E. coli* at the point $\mathbf{x}$ (defined by $s$ or $\sigma$) on the interface at time $t$. This expression assumes that the expansion velocity is reduced in linear proportion to the local *E. coli* concentration $c$ with proportionality coefficient $\alpha$ and is also subject to the linear surface tension with coefficient $\gamma$. Then, the model (S6),(S7) can be rewritten as

$$\dot{\kappa} = -\left(\frac{\partial^2}{\partial s^2} + \kappa^2\right)(F_0 - \gamma\kappa - \alpha c_0/g) \tag{S9}$$

$$\dot{g} = 2g\kappa(F_0 - \gamma\kappa - \alpha c_0/g) \tag{S10}$$

We can perform a linear stability analysis of a flat interface ($\kappa = 0, g = 1$) by substituting ansatz

$$\kappa = Ke^{iks+\lambda t} \tag{S11}$$

$$g = 1 + Ge^{iks+\lambda t} \tag{S12}$$

in Eqs. (S9), (S10). The Jacobian of the linearized system reads

$$J = \begin{bmatrix} -\gamma k^2 & \alpha c_0 k^2 \\ 2(F_0 - \alpha c_0) & 0 \end{bmatrix}. \tag{S13}$$

For positive $\gamma, \alpha$ and $F_0 > \alpha c_0$ (the latter condition means that the colony with smooth interface is expanding), one of the two eigenvalues of this Jacobian is always positive. At small wavenumbers $k$, it increases linearly with $k$,

$$\lambda = \sqrt{2\alpha c_0(F_0 - \alpha c_0)}k \tag{S14}$$

and for large $k$ it reaches the maximum value

$$\lambda_m = \frac{2\alpha c_0(F_0 - \alpha c_0)}{\gamma} \tag{S15}$$

Since the growth rate is positive for all values of $k$, this instability may lead to singularities in curvature (cusps). This is indeed what is found in numerical simulations of the discrete analog of this model (see the next section). These singularities correspond to the origins of "branches" of *E. coli* that the interface leaves behind during the flower pattern growth.

#### Flexible-chain interface model

The interface dynamics beyond linear instability stage can be analyzed numerically. Unfortunately, it is difficult to implement self-avoidance of the interface in the framework of the continuum model described in the previous section. Thus, we implemented a discrete flexible-chain model that is analogous to the continuum model described above but contains additional interaction terms between the nodes that prevent self-intersection of the chain. Specifically, we represent the interface as a closed chain of $N$ nodes with coordinates $\mathbf{x}_i, i = 1, ..., N$. Let us introduce the vectors connecting node $i-1$ to node $i$ (we assume that node 0 is the same as node N): $\mathbf{\Delta}_i = \mathbf{x}_i - \mathbf{x}_{i-1}$. Each node is driven by the "expansion force" $F_0$ that acts along the unit vector $\hat{\mathbf{n}}_i$ that is directed outwards along the bisectrix of two adjacent edges, $\mathbf{\Delta}_i$ and $\mathbf{\Delta}_{i+1}$. It is counteracted by the "friction" force that is directed along $-\hat{\mathbf{n}}_i$ and is proportional to the local density of *E. coli* $g_i$ associated with node $i$ and by the surface tension force that is proportional to the local curvature of the interface $\kappa_i$. In addition, we introduce repulsion forces between all nodes and edges that prevent the interface from self-intersecting. The equation of motion in the overdamped limit can be written as follows:

$$\frac{d\mathbf{x}_i}{dt} = \hat{\mathbf{n}}_i(F_0 - \alpha g_i - \gamma\kappa_i) + \sum_{j\neq i} \mathbf{f}_{ij}^{nn} + \sum_{j\neq i} \mathbf{f}_{ij}^{ne} \tag{S16}$$

The discrete analog of the local curvature at node $i$ is defined as follows,

$$\kappa_i = \left| \frac{\mathbf{\Delta}_{i+1}}{\Delta_{i+1}} - \frac{\mathbf{\Delta}_i}{\Delta_i} \right| \frac{2}{\Delta_i + \Delta_{i+1}} \tag{S17}$$

where $\Delta_i = |\mathbf{\Delta}_i|$.

We assume that each node carries the fixed "amount" of *E. coli* $g$, and the local concentration of *E. coli* $g_i$ is defined as the average amount of $g$ per unit length of the interface. In the simplest case, it can be computed as $g/L_i$ where $L_i$ is the half-sum of lengths of two edges attached to node $i$, $L_i = \frac{1}{2}(\Delta_i + \Delta_{i+1})$, however in simulations we typically used longer averaging over 2 adjacent edges on both sides,

$$g_i = \frac{2(2K+1)g}{\sum_{j=-K}^{K}[\Delta_{i+j} + \Delta_{i+1+j}]} \tag{S18}$$

with $K = 2$.

The last two terms in the r.h.s. of Eq.(S16) represents the vector sum of possible repulsive forces acting on the node $i$ from other nodes ($\mathbf{f}_{ij}^{nn}$) or edges ($\mathbf{f}_{ij}^{ne}$) of the chain. The node-node force acts along the vector connecting nodes $i$ and $j$, $\mathbf{x}_i - \mathbf{x}_j$. We assume that the node-edge force acts perpendicular to the orientation of the $j$-th link, $\Delta_j$. We assume that the node-node force $\mathbf{f}_{ij}^{nn}$ is zero if $d_{ij}^{nn} = |\mathbf{x}_i - \mathbf{x}_j| > d_0$ and varies as $F_m(1 - d_{ij}^{nn}/d_0)^4$ for $d_{ij}^{nn} < d_0$ with large $F_m \ll F_0$. Similarly, the node-edge force $\mathbf{f}_{ij}^{ne}$ is zero if the distance between the node $i$ and the edge $j$, $d_{ij}^{ne} > d_0$ and varies as $F_m(1 - d_{ij}^{ne}/d_0)^4$ for $d_{ij}^{ne} < d_0$.

#### Parameters

We used parameters below unless specified otherwise.

| $F_0$ | $\alpha$ | $\gamma$ | $F_m$ | $d_0$ | $g_0$ | $N$ | $dt$ |
|---|---|---|---|---|---|---|---|
| 1 | 0.5 | $10^{-8}$ | 0.1 | 0.01 | 1 | 512 | 0.001 |

### Phase-field model

#### Model description

In this more elaborate 2D model of a two-strain biofilm, we consider it as a growing mass of compressible fluid. A convenient way to describe a compact expanding biofilm is to use a phase-field approach where the phase $\phi$ changes smoothly from 0 outside the colony to 1 inside. The evolution of phase field $\phi$ is given by the equation:

$$\frac{\partial\phi}{\partial t} = -\mathbf{u}\cdot\nabla\phi + \Gamma(\epsilon\nabla^2\phi - G'(\phi)/\epsilon + \kappa\epsilon|\nabla\phi|) \tag{S19}$$

where $\mathbf{u}$ is the velocity field, $\kappa = -\nabla\cdot(\nabla\phi/|\nabla\phi|)$ is the local interface curvature, and $\epsilon$ characterizes the interface width. The term $G(\phi) = 18\phi^2(1-\phi)^2$ is included to force the bistable dynamics of $\phi$ field with two stable fixed points at 0 and 1. The dynamics of the *A. baylyi* cells density $\rho_A$ within the biofilm is described by

$$\frac{\partial(\phi\rho_A)}{\partial t} + \nabla\cdot(\phi\rho_A\mathbf{u}) = \nabla\cdot(\phi D_A\nabla\rho_A) + \alpha_A\phi\rho_A(1-\rho_A-\rho_E) \tag{S20}$$

The second term in the left-hand side is the advection term while the two terms in the right-hand side are diffusion and growth terms respectively. $D_A$ and $\alpha_A$ are the diffusion constant and growth rate of *A. baylyi* respectively. The growth term follows logistic form and we assume that the growth can be saturated when the total density of *A. baylyi* ($\rho_A$) and *E. coli* ($\rho_E$) reaches 1.

Similarly, the dynamics for *E. coli* cells density $\rho_E$ is described by

$$\frac{\partial(\phi\rho_E)}{\partial t} + \nabla\cdot(\phi\rho_E\mathbf{u}) = \nabla\cdot(\phi D_E\nabla\rho_E) + \alpha_E\phi\rho_E(1-\rho_A-\rho_E) \tag{S21}$$

where $D_E$ and $\alpha_E$ are the diffusion rate and growth rate of *E. coli.* Note that the advection of the phase field and both cell densities is provided by the same velocity field $\mathbf{u}$.

The velocity field could be determined by the overdamped Stokes equation:

$$\nabla\cdot[\nu(\phi)(\nabla\mathbf{u}+\nabla\mathbf{u}^{\mathbf{T}})] + \nabla\cdot(\chi\sigma_A) - [\xi+\beta f(\rho_E)\phi]\mathbf{u} = 0 \tag{S22}$$

where $\nu(\phi) = \nu_0\phi$ is the viscosity, $\sigma_A = -\eta\phi\rho_A\mathbf{I}$ is the stress provided by motile *A. baylyi* cells ($\mathbf{I}$ is the identity matrix). $\chi$ is a random number uniformly distributed between $1 \pm 0.3$, which adds noise to the stress driven by *A. baylyi.* Because pure *E. coli* colony expands very slowly and pure *A. baylyi* colony expands fast, we assume that the stress provided by *E. coli* is negligible compared to *A. baylyi.* Our experiments with mixtures of *E. coli* and *A. baylyi* show that regions where there are more *E. coli* move outward more slowly, so we assume that *E. coli* cells provide friction to prevent colony from expanding fast. This is described by the last term in which $\xi$ is the basal friction constant and $f(\rho_E) = \rho_E$ determines how the friction is modulated by *E. coli* cells. Here we assume it is simply proportional to $\rho_E$.

#### Numerical algorithm

The numerical algorithm is similar to [3]. We aim to solve Eqns. S19-S22 with uniform spatial grid sizes $\Delta x, \Delta y$ and fixed time step $\Delta t$ from initial conditions $\phi^0, \rho_A^0, \rho_E^0, \mathbf{u}^0$. The system variables at time $t = n\Delta t$ are denoted as $\phi^n, \rho_A^n, \rho_E^n, \mathbf{u}^n$.

We first solve Eq. S19 by forward Euler scheme:

$$\phi^{n+1} = \phi^n - \Delta t\mathbf{u}^n\cdot\nabla\phi^n + \Delta t\Gamma[\epsilon\nabla^2\phi^n - G'(\phi^n)/\epsilon + \epsilon\kappa^n|\nabla\phi^n|]$$

with $\kappa^n$ calculated by $\kappa^n = -\nabla\cdot(\nabla\phi^n/|\nabla\phi^n|)$ when $|\nabla\phi^n| > 0.01$, and set to 0 otherwise.

The reaction-diffusion-advection equations for $\rho_A$ and $\rho_E$ are discretized using the forward Euler scheme:

$$\phi^n\frac{\rho^{n+1}-\rho^n}{\Delta t} + \frac{\phi^{n+1}-\phi^n}{\Delta t}\rho^n = \text{Advection+Diffusion+Reaction} \tag{S23}$$

where $\phi^{n+1}$ is obtained from the above step, and $\rho^{n+1}$ is only updated when $\phi^n > 10^{-4}$. The advection term is calculated by

$$\begin{aligned}[\nabla\cdot(\phi^n\rho^n\mathbf{u}^n)]_{ij} &= (\phi^n_{i+1/2,j}\rho^n_{i+1/2,j}u^n_{i+1/2,j} - \phi^n_{i-1/2,j}\rho^n_{i-1/2,j}u^n_{i-1/2,j})/\Delta x \\ &\quad +(\phi^n_{i,j+1/2}\rho^n_{i,j+1/2}v^n_{i,j+1/2} - \phi^n_{i,j-1/2}\rho^n_{i,j-1/2}v^n_{i,j-1/2})/\Delta y\end{aligned}$$

and for the diffusion term

$$\begin{aligned}[\nabla\cdot(\phi^n D\nabla\rho^n)]_{ij} &= D[\phi_{i+1/2,j}\frac{\rho_{i+1,j}-\rho_{i,j}}{\Delta x} - \phi_{i-1/2,j}\frac{\rho_{i,j}-\rho_{i-1,j}}{\Delta x}]/\Delta x \\ &\quad +D[\phi_{i,j+1/2}\frac{\rho_{i,j+1}-\rho_{i,j}}{\Delta y} - \phi_{i,j-1/2}\frac{\rho_{i,j}-\rho_{i,j-1}}{\Delta y}]/\Delta y\end{aligned}$$

where $\mathbf{u} = (u, v)$, $\phi_{i\pm1/2,j} = (\phi_{i\pm1,j} + \phi_{i,j})/2$, $\phi_{i,j\pm1/2} = (\phi_{i,j\pm1} + \phi_{i,j})/2$, and we used the same definitions for $\rho$, $u$ and $v$ between collocation points. Then we can calculate $\rho^{n+1}$ from Eqn. S23.

The Stokes equation Eq. S22 is integrated by the semi-implicit Fourier spectral method (to stabilize the scheme, we subtract the term $\nu_0\phi_0\nabla^2\mathbf{u}$ from both sides of Stokes equation with large constant $\phi_0$, e.g. $\phi_0 = 100$):

$$\xi\mathbf{u} - \nu_0\phi_0\nabla^2\mathbf{u} = \nu_0\nabla\cdot[\phi\nabla\mathbf{u}^T + (\phi-\phi_0)\nabla\mathbf{u}] + \nabla\cdot(\chi\sigma_A) - \beta f(\rho_E)\phi\mathbf{u}$$

To obtain $\mathbf{u}^{n+1}$, we set $\mathbf{u}_0^{n+1} = \mathbf{u}^n$ and solve the equation below iteratively using spectral Fourier method:

$$\xi\mathbf{u}_{k+1}^{n+1} - \nu_0\phi_0\nabla^2\mathbf{u}_{k+1}^{n+1} = \nu_0\nabla\cdot[\phi^{n+1}\nabla\mathbf{u}_k^{T,n+1} + (\phi^{n+1}-\phi_0)\nabla\mathbf{u}_k^{n+1}] + \nabla\cdot(\chi\sigma_A)^{n+1} - \beta f(\rho_E^{n+1})\phi^{n+1}\mathbf{u}_k^{n+1}$$

where $k = 0, 1, 2, \cdots$ are iteration steps. In simulations, we constrain the error by iterating the above process until

$$\max|\mathbf{u}_k^{n+1} - \mathbf{u}_{k-1}^{n+1}| < 0.01\max|\mathbf{u}_k^{n+1}|$$

or until $k_{max} = 200$, and the final $\mathbf{u}^{n+1} = \mathbf{u}_m^{n+1}$.

#### Parameters

We used parameters below unless specified otherwise.

| $\Gamma$ | $\epsilon$ | $D_A$ | $\alpha_A$ | $D_E$ | $\alpha_E$ | $\nu_0$ | $\eta$ | $\xi$ | $\beta$ | $\Delta x$ | $\Delta y$ | $\Delta t$ |
|---|---|---|---|---|---|---|---|---|---|---|---|---|
| 0.4 | 8.1 | 6.1 | 2 | 0.1 | 2.2 | 9 | 75 | 1 | 18 | 0.4 | 0.4 | 0.0001 |

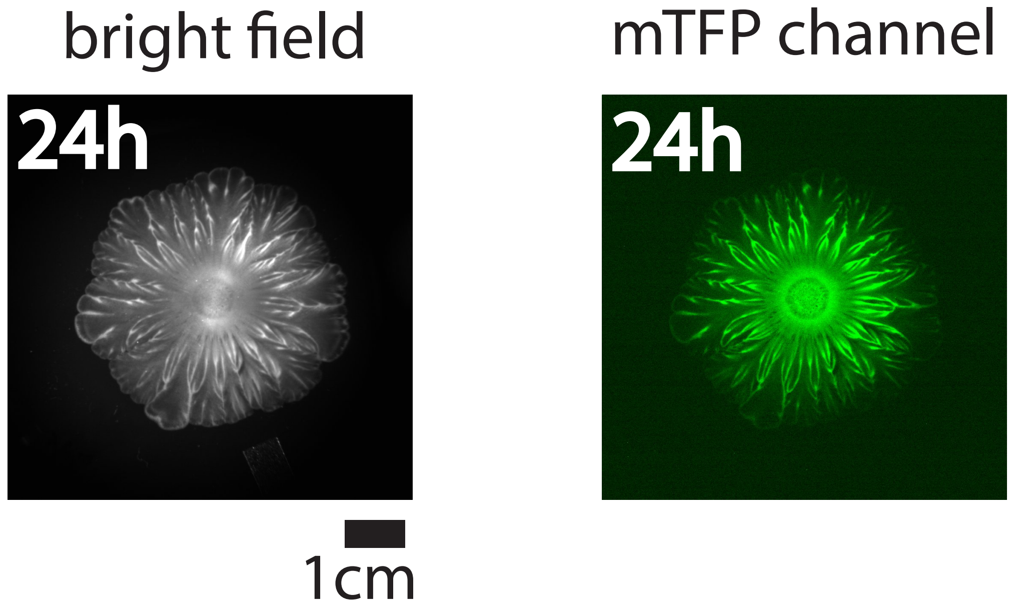


Figure S1: Bright-field image (left) and mTFP channel image (right) for the flower-like pattern after 24 hours of growth under milliscope.

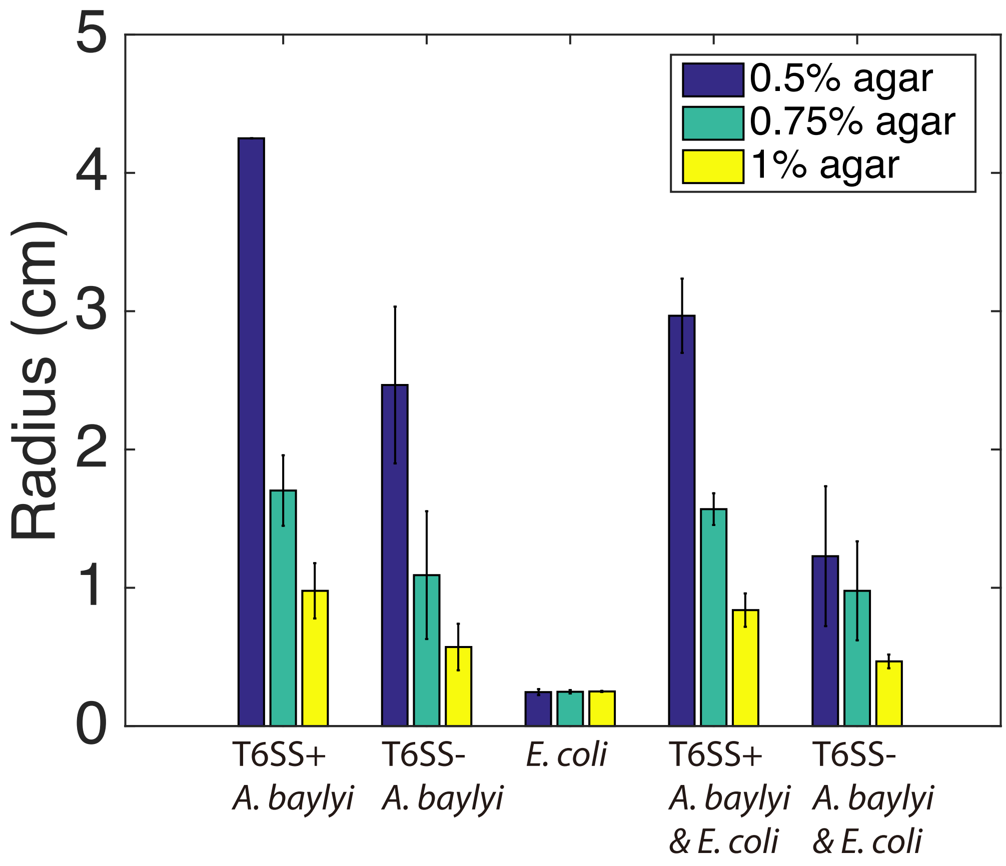


Figure S2: Average colony radii after 16 hours of growth in 37°C for pure T6SS$^+$ *A. baylyi*, pure T6SS$^-$ *A. baylyi*, pure *E. coli*, mixture of T6SS$^+$ *A. baylyi* and *E. coli* with 1:1 initial density ratio, mixture of T6SS$^-$ *A. baylyi* and *E. coli* with 1:1 initial density ratio with different agar concentrations (10 mL LB agar). For pure T6SS$^+$ *A. baylyi* on LB agar (0.5% agar) plate, after 16 hours, the colony already reached the edge of the plate, so the radius of the plate is shown here. For each combination, experiments were run in triplicate.

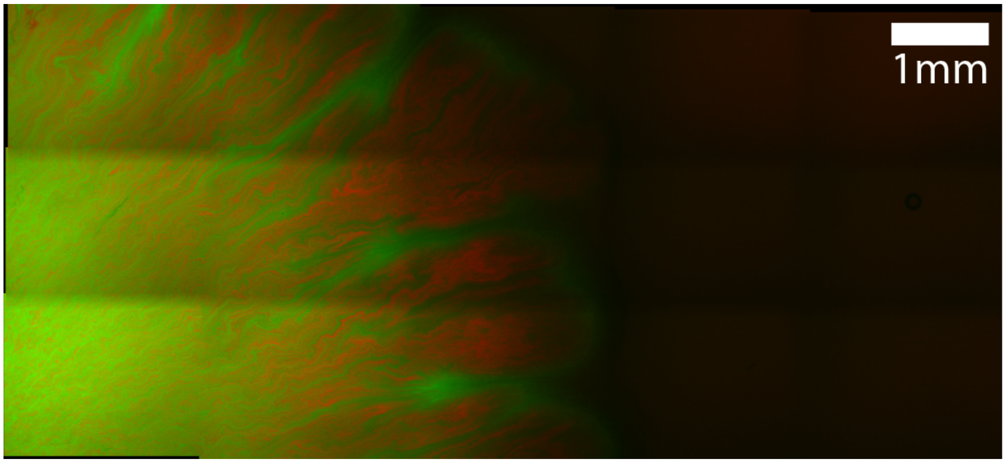


Figure S3: Microscope image of mixture of *E. coli* and T6SS$^-$ *A. baylyi* on agar surface. Red color shows *A. baylyi* (mCherry channel) while green color shows *E. coli* (mTFP channel).

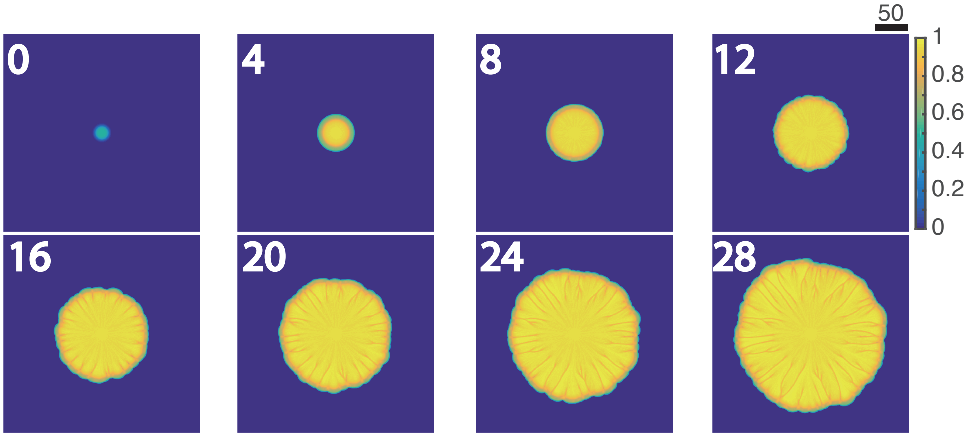


Figure S4: Several snapshots of *A. baylyi* density during the growth of a mixed colony in a phase-field model simulation.

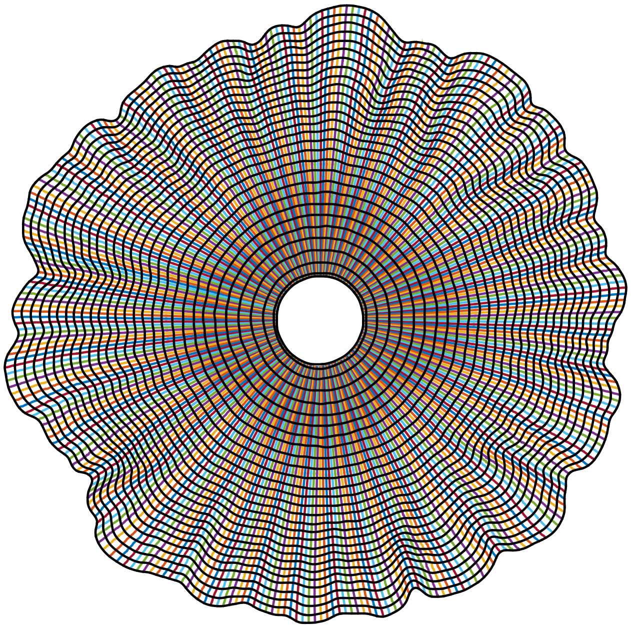

Figure S5: Examples of the tracked colony boundary and traces of 300 virtual nodes on the colony boundary. Black curves show the splines interpolated from the positions of 300 nodes at each time point. Different curves with colors show the traces of each node.

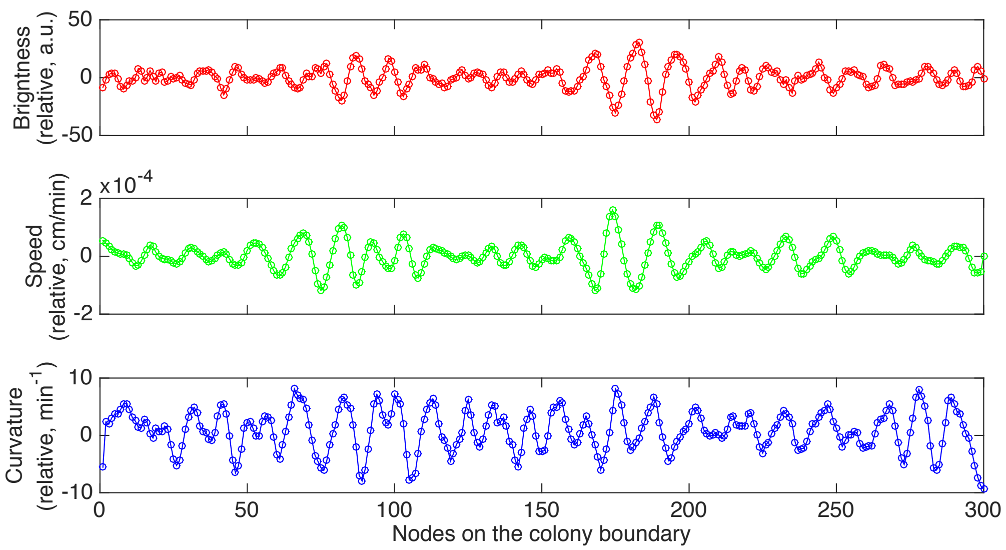


Figure S6: An example of the relative brightness, speed and local cuvature for all 300 nodes after 10 hours of colony growth.

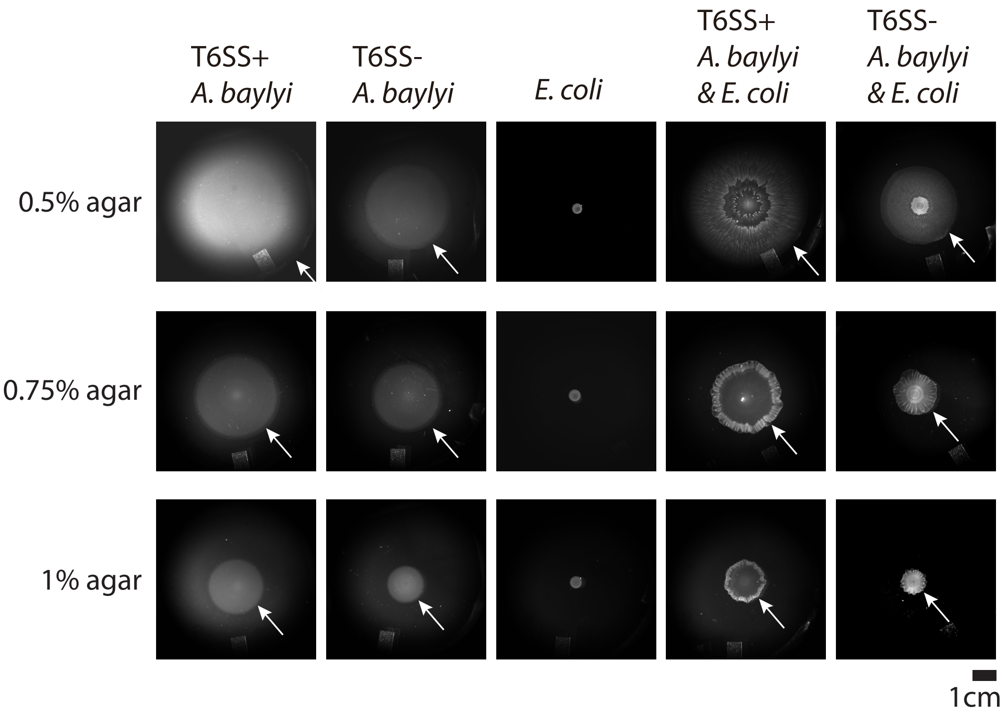


Figure S7: Examples of the colonies for all combinations of *E. coli* and *A. baylyi* with different agar concentrations after 16 hours of growth on 10 mL LB agar. When *A. baylyi* and *E. coli* were mixed, the initial seeding density ratio was 1:1.

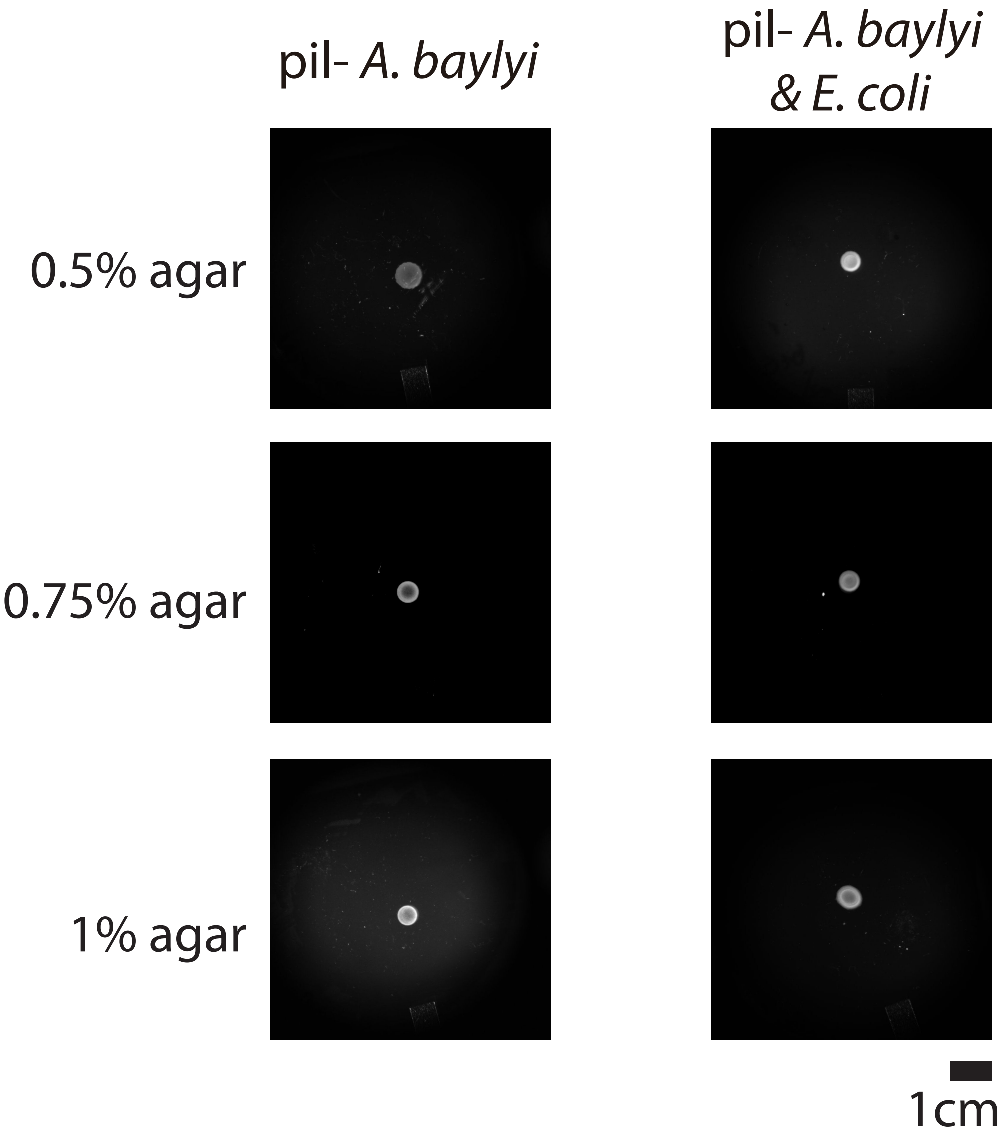


Figure S8: Examples of the colonies for pure pil$^{-}$ T6SS$^{+}$ *A. baylyi*, mixture of pil$^{-}$ T6SS$^{+}$ *A. baylyi* and *E. coli* with initial seeding density ratio 1:1 with different agar concentrations after 16 hours of growth on 10 mL LB agar.

#### Supplementary Movies

**Movie S1** Formation of flower-like patterns in the mixture of T6SS$^+$*A. baylyi* and *E. coli* under milliscope. Initial A:E density ratio was 1:10 and the cells grew on 10 mL LB agar (0.5% agar).
**Movie S2** Development of branches in a growing flower-like pattern under microscope (4x magnification). Initial A:E density ratio was 1:1 and the cells grew on 10 mL LB agar (1% agar).
**Movie S3** A sample simulation of the discrete biofilm interface model.
**Movie S4** A pattern forming as a "fossil record" of node colors (corresponding to *E. coli*density on the interface) in the discrete interface model simulation of Movie S3.
**Movie S5** A sample simulation of the phase-field model of two-species colony growth.
**Movie S6** When T6SS$^+$ *A. baylyi* and *E. coli* we inoculated separately on 10 mL LB agar (0.75% agar), the flower pattern formed only in a segment.
**Movie S7** Segmentation and tracking of the boundary of the growing colony from Movie S1.
